# RefSoil: A reference database of soil microbial genomes

**DOI:** 10.1101/053397

**Authors:** Jinlyung Choi, Fan Yang, Ramunas Stepanauskas, Erick Cardenas, Aaron Garoutte, Ryan Williams, Jared Flater, James M Tiedje, Kirsten S. Hofmockel, Brian Gelder, Adina Howe

## Abstract

A database of curated genomes is needed to better assess soil microbial communities and their processes associated with differing land management and environmental impacts. Interpreting soil metagenomic datasets with existing sequence databases is challenging because these datasets are biased towards medical and biotechnology research and can result in misleading annotations. We have curated a database of 922 genomes of soil-associated organisms (888 bacteria and 34 archaea). Using this database, we evaluated phyla and functions that are enriched in soils as well as those that may be underrepresented in RefSoil. Our comparison of RefSoil to soil amplicon datasets allowed us to identify targets that if cultured or sequenced would significantly increase the biodiversity represented within RefSoil. To demonstrate the opportunities to access these underrepresented targets, we employed single cell genomics in a pilot experiment to sequence 14 genomes. This effort demonstrates the value of RefSoil in the guidance of future research efforts and the capability of single cell genomics as a practical means to fill the existing genomic data gaps.

## Introduction

Microbial populations in the soil impact the structure and fertility of our soils by contributing to nutrient availability, soil stability, plant productivity, and protection against disease and pathogens. Soil microbiology has a rich history of isolating and characterizing representatives from the soil to assess their impacts on soil health and stability. However, despite significant efforts to isolate microbes from the soil, we have accessed only a small fraction of its biodiversity, with estimates of less than 1 to 50 percent of species, even with novel isolation techniques (Van H T Pham and Kim, 2012; Vartoukian *et al.*, 2010; Schoenborn *et al.*, 2004; Burmølle *et al.*, 2009; Janssen *et al.*, 2002; Ling *et al.*, 2015). With advances in DNA sequencing technologies, we can now directly interrogate soil metagenomes, allowing for the characterization of soil communities without the need to first cultivate isolates. However, our ability to annotate and characterize the retrieved genes is dependent on the availability of informative reference gene or genome databases. The largest resource of full genome reference sequences for annotating genes is the NCBI RefSeq database(Tatusova *et al.*, 2013), which contains a total of 57,993 genomes (Release 74). While broadly useful, this database is not representative of the large majority of soil microbiomes, and the majority of genes in previously published soil metagenomes (65-90%) cannot be annotated against known genes(Delmont *et al.*, 2012; Fierer *et al.*, 2012). Further, contributions to the NCBI databases have largely originated from human health and biotechnology research efforts that can mislead annotations of genes originating from soil microbiomes (e.g., annotations that are clearly not compatible with life in soil).

Soil microbiologists are not the first to face the problem of a limited reference database. The NIH Human Microbiome Project (HMP) recognized the critical need for a well-curated reference genome dataset and developed a reference catalog of 3,000 genomes that were isolated and sequenced from human-associated microbial populations(Huttenhower *et al.*, 2012). This publicly available reference set of microbial isolates and their genomic sequences aids in the analysis of human microbiome sequencing data(Segata *et al.*, 2012; Wu *et al.*, 2009) and also provides strains for which isolates (both culture collections and nucleic acids) are available as resources for experiments. Following this successful model of the HMP, we have developed a curated database of reference genome sequences that originate from soil. This database, called RefSoil, represents the current state of knowledge of sequenced soil genomes. In addition to helping us to understand known soil microbiology, RefSoil aids in the identification of knowledge gaps that need to be filled to improve our understanding of soil biodiversity. We discuss the importance the RefSoil database and demonstrate exciting opportunities to expand this database and its applications in the future.

## Materials and Methods

### Creation of RefSoil Database

In order to create a soil-specific reference genome dataset, we targeted genome sequences based on evidence that isolates had origins in soil systems. A total of 6,646 complete genomes were obtained from the Genomes OnLine Database (GOLD, https://gold.jgi.doe.gov, October 9th, 2014); the GOLD database was chosen because of the availability of metadata, particularly environmental origin, related to genome sequences. These genomes were further selected based on soil-association with the following criteria: (1) Within the GOLD database, organism information and organism metadata (known habitats, ecosystem category, and ecosystem type) was required to identify organism as originating in soil-associated categories. Organisms from marine and deep sea environments were excluded, though these organisms often were identified as soil-associated. Additionally, obligate host-associated pathogens and extremophiles were excluded, as these organisms are unlikely to be present in the absence of their host or in representative soil samples. We considered these organisms to often be under very strong selective pressures that can lead to reduced genomes or high rates of recombination that are difficult to assess soil-specific trends. (2) For organisms that lacked appropriate metadata in GOLD, a Google Custom Search was used to query all available websites for the organism name and soil-related terms (rhizosphere, soil, sand, mud, or nodule). Priorities were placed on the following websites: http://www.ncbi.nlm.nih.gov/pubmed/, http://www.straininfo.net/, https://microbewiki.kenyon.edu/index.php/MicrobeWiki,https://en.wikipedia.org/, and https://scholar.google.com. Genomes with an association with at least one webpage containing search query phrases were included in RefSoil. This approach was tested with known soil-associated organisms, verifying that it would reliably predict soil association for these organisms. Within resulting genomes, duplicated chromosomes and strains were removed. If multiple genome accession numbers were associated with a single strain, the most recent genome sequence was chosen. NCBI Genbank annotation files associated with each strain were obtained on 2/18/16. The genomes contained within RefSoil are provided in Supplementary Table 1.

### Characterization of organisms in RefSoil

All RefSoil genome sequences were associated with their NCBI RefSeq accession ID and obtained from NCBI. Using NCBI Genbank CDS annotations, 16s rRNA gene sequences were identified. If multiple 16S rRNA gene sequences were present, the first sequence from the first chromosome (by NCBI index) was selected as representative for the genome. Three genomes lacked annotations of a 16S rRNA gene and a 16S rRNA gene HMM model was used and identified an additional 16S rRNA sequence(Guo *et al.*, 2015). In total, 886 16S rRNA gene sequences were used to build a phylogenetic tree for bacterial genomes within RefSoil (Fig. 1). All bacterial 16S rRNA gene sequences were aligned using RDPs bacterial model (Infernal 1.1.1 (Nawrocki and Eddy, 2013)), and a Maximum-likelihood phylogenetic tree was constructed based on the Jukes-Cantor model by Fasttree (Price *et al.*, 2010) and visualized with Graphlan (R 3.2.2, version 0.9.7) (Fig. 1). The scripts for these approaches are publicly available on https://github.com/germs-lab/ref_soil. RefSoil genomes were extracted from corresponding genomes contained within NCBI RefSeq (release 74). Annotations of taxonomy for RefSoil and RefSeq genes were obtained from NCBI (February 19, 2016) (Supplementary Fig. 1). RefSoil genomes were submitted to RAST and genes were annotated using SEED subsystem ontology (Supplementary Table 1). Among 1,811,233 unique genes, approximately 39.1% of them were assigned to multiple SEED subsystem level 1 categories. For the purpose of this study, we included all annotations at subsystem level 1 for each unique gene, which expanded the total gene counts to 2,619,643. The percent abundance of genes from each phylum in RefSoil database was also adjusted accordingly by including the abundance of genes that were assigned to multiple subsystem level 1 categories (Fig. 2 and Supplement Table 2). The number of phyla identified in each subsystem level 1 categories was summarized in Supplemental Table 3.

**Figure 1.**
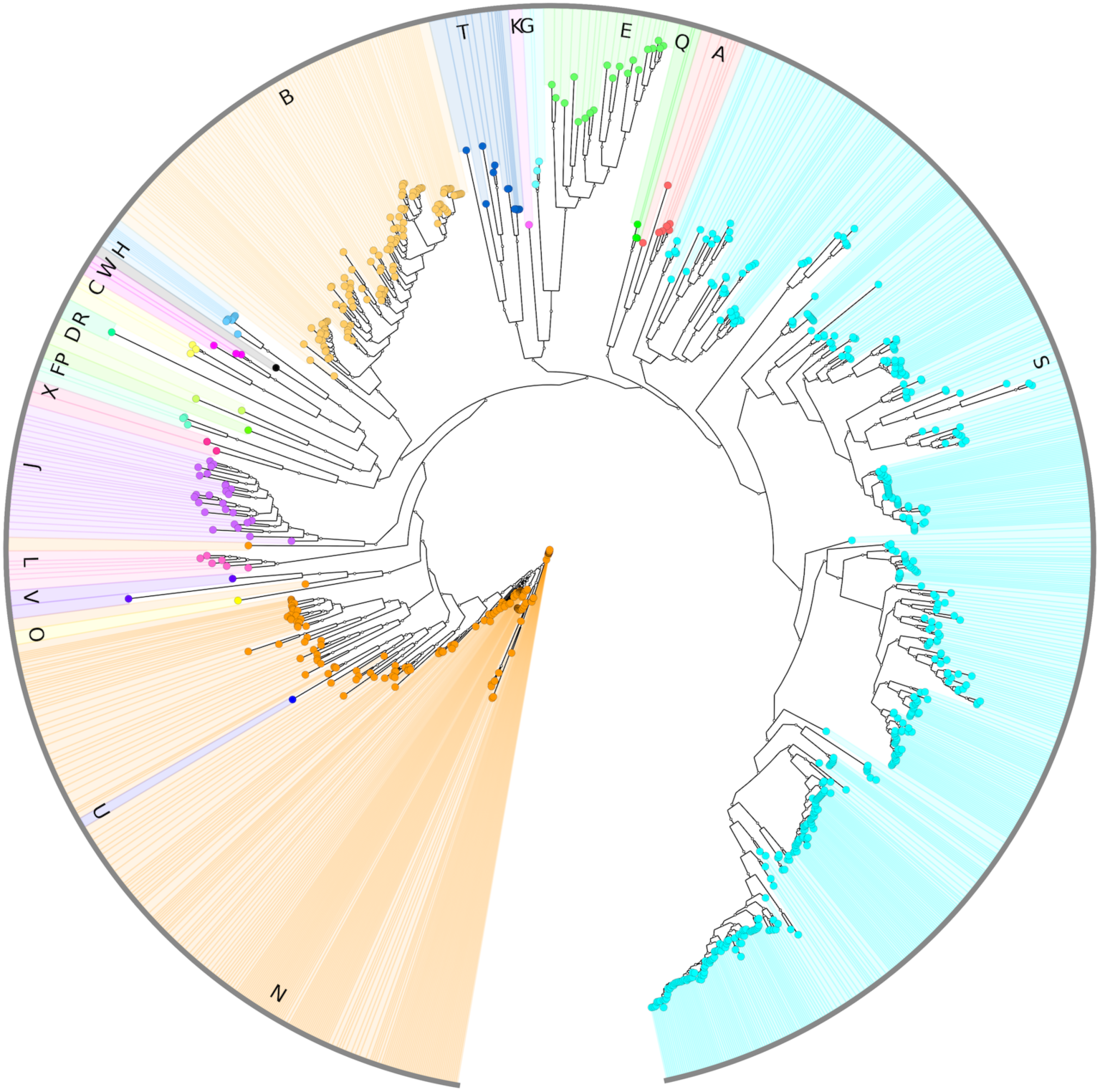
Phylogenetic tree of RefSoil. Phylogenetic tree of aligned 16S rRNA genes originating from RefSoil bacterial genomes. A: Acidobacteria, B: Actinobacteria, C: Aquificae, D: Armatimonadetes, E: Bacteroidetes, F: Chlamydiae, G: Chlorobi, H: Chloroflexi, J: Cyanobacteria, K: Deferribacteres, L: Deinococcus-Thermus, N: Firmicutes, O: Fusobacteria, P: Gemmatimonadetes, Q: Nitrospirae, R: Planctomycetes, S: Proteobacteria, T: Spirochaetes, U: Synergistetes, V: Tenericutes, W: Thermotogae, X: Verrucomicrobia.

**Figure 2.**
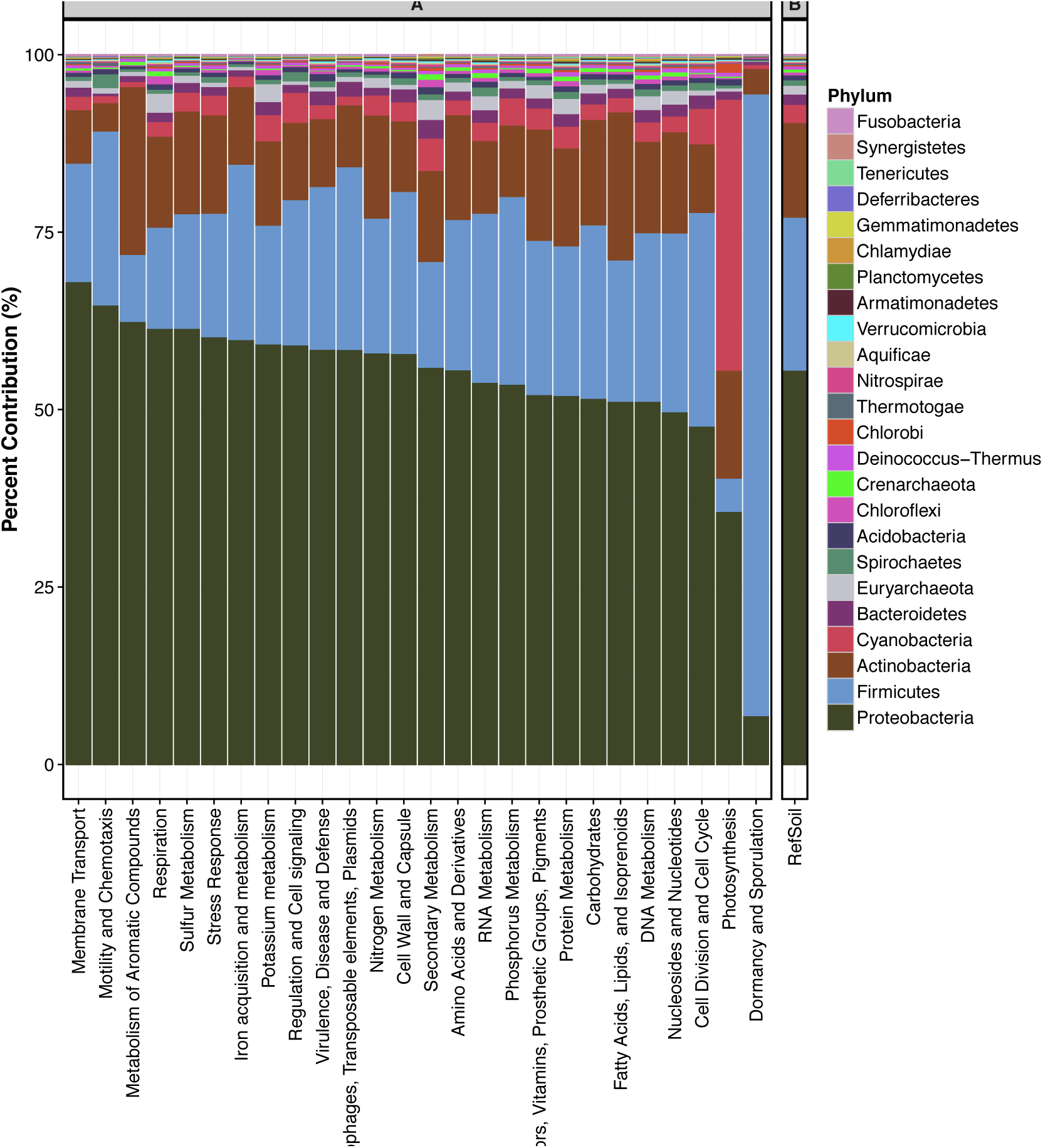
Functional analysis. The distribution of phylogenetic origins associated with RefSoil functional subsystems as annotated by RAST (panel A); the overall phylogenetic distribution of all genes in RefSoil (panel B).

### EMP databases used in this study

A total of 15,481 amplicon datasets that form the EMP dataset were available to compare environmental soil amplicons to RefSoil (total 5,594,412 OTUs, clustered at 97%)(Rideout *et al.*, 2014). Soil samples were selected based on their association with soil metadata resulting in a total of 2,476,795 OTUs from 3035 samples. A total of 2,158 unique taxonomy assignments were identified for EMP OTUs using the RDP Classifier (Wang *et al.*, 2007) and used to construct a phylogenetic tree with the Graphlan package (R 3.2.2, version 0.9.7) (Fig. 3). Abundances for each taxonomic assignment were calculated as the sum of abundance of OTUs associated with that taxonomy.

**Figure 3.**
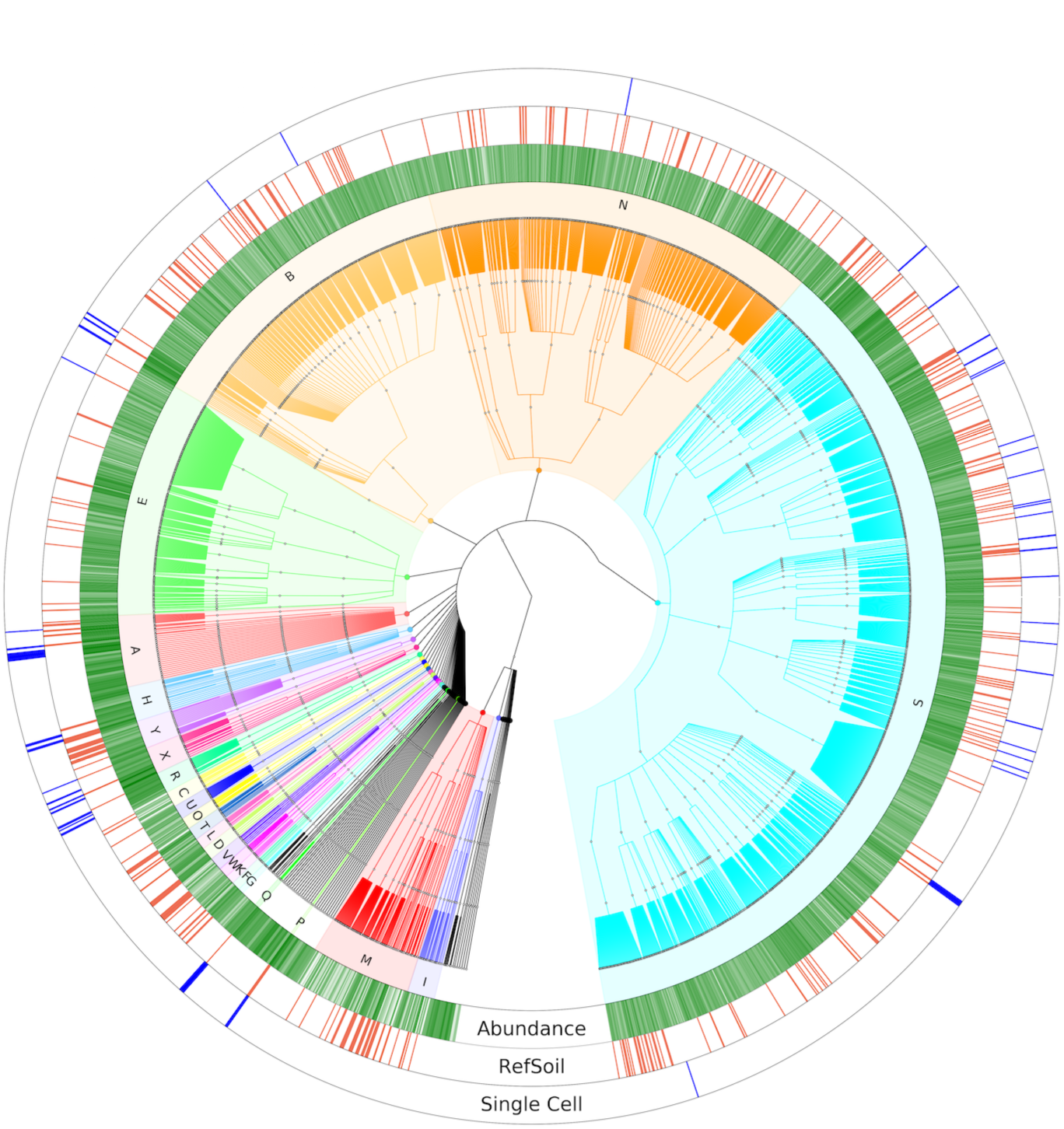
Phylogenetic tree of EMP OTUs clustered by taxonomy. Ring I (green) represents the cumulative log-scaled abundance of OTUs in EMP soil samples. Ring II (red) represents EMP OTUs that share greater than 97% gene similarity (to RefSoil 16S rRNA genes; ring III (blue) indicates that these 16S rRNA genes shared similarity to sorted cells that were selected for single cell genomics. A: Acidobacteria, B: Actinobacteria, C: Aquificae, D: Armatimonadetes, E: Bacteroidetes, F: Chlamydiae, G: Chlorobi, H: Chloroflexi, I: Crenarchaeota, K: Deferribacteres, L: Deinococcus-Thermus, M: Euryarchaeota, N: Firmicutes, O: Fusobacteria, P: Gemmatimonadetes, Q: Nitrospirae, R: Planctomycetes, S: Proteobacteria, T: Spirochaetes, U: Synergistetes, V: Tenericutes, W: Thermotogae, X: Verrucomicrobia, Y: Cyanobacteria/Chloroplast.

To evaluate the presence of RefSoil genomes in EMP, EMP and RefSoil 16S rRNA gene amplicons were compared by alignment with BLAST, requiring an alignment with greater than 97% similarity, a minimum alignment of 72 bp, and E-value ≤ 1e-5. If multiple hits were aligned, the hit with the lowest E-value was selected as representative. Using these criteria, a total of 53,538 EMP OTUs were associated RefSoil 16s rRNA genes (1.4% of total EMP OTUs; 10.2% of all EMP amplicons).

Soil order for EMP samples located in the United States (1817 samples) was obtained based on GPS location associated with sample metadata; locations were only considered valid if the coordinates entered were actually located in the United States (1627 samples). Valid GPS points were then located within a soil order by querying the USDA NRCS Global Soil Regions map (Reich) (Accessed on January 26, 2016) for soil order plus a Rock/Sand/Ice category using ArcGIS 10.3.1. RefSoil representatives for EMP OTUs were determined by similarity as described above (Supplementary Table 4).

### Most-wanted soil OTUs

The selection of the most wanted OTUs for RefSoil based on EMP-associated amplicons was based on presence in EMP samples, using the criteria of EMP-RefSoil similarity as described above, and abundance (number of amplicons associated with the OTU in EMP samples). Candidate OTUs were ranked based on its observed frequency in all EMP samples and abundance in EMP amplicons (Top 100 shown in Supplementary Table 5 and 6). Taxonomy was assigned by the RDP classifier (Wang *et al.*, 2007). The 21 OTUs present in both the most abundance and the most frequent are listed in the (Table 1.

**Table 1.** RefSoils’s most wanted OTUs. RefSoils most wanted OTUs based on observed frequency and abundance in EMP soil samples. Taxonomy for OTUs are assigned by RDP classifier(Wang *et al.*, 2007).

| OTU ID in<br>(Rideout et<br>al., 2014) | Closest Match in RDP Classifier |  | RDP Classifier<br>Similarity Score | Abundance<br>(total<br>amplicons) | Number<br>of<br>Samples |
| --- | --- | --- | --- | --- | --- |
|  | Phylum | Class |  |  |  |
| 4457032 | Verrucomicrobia | Spartobacteria | 1 | 8,007,453 | 2,312 |
| 4471583 | Verrucomicrobia | Spartobacteria | 0.92 | 2,937,242 | 1,935 |
| 101868 | Verrucomicrobia | Spartobacteria | 1 | 1,828,706 | 2,151 |
| 559213 | Firmicutes | Bacilli | 1 | 1,606,757 | 2,410 |
| 1105039 | Verrucomicrobia | Spartobacteria | 1 | 1,295,847 | 2,102 |
| 807954 | Bacteroidetes | Sphingobacteriia | 0.98 | 875,988 | 2,546 |
| 4342972 | Verrucomicrobia | Spartobacteria | 0.97 | 750,557 | 2,386 |
| 4423681 | Gemmatimonadetes | Gemmatimonadetes | 0.25 | 689,209 | 1,954 |
| 1109646 | Verrucomicrobia | Spartobacteria | 1 | 554,748 | 2,012 |
| 610188 | Acidobacteria | Acidobacteria_Gp6 | 0.99 | 553,476 | 2,694 |
| 4373456 | Acidobacteria | Acidobacteria_Gp4 | 1 | 397,255 | 2,150 |
| 4341176 | Verrucomicrobia | Subdivision3 | 0.99 | 383,900 | 2,098 |
| 720217 | Proteobacteria | Deltaproteobacteria | 0.74 | 345,621 | 2,073 |
| 4314933 | Acidobacteria | Acidobacteria_Gp1 | 0.8 | 333,428 | 1,974 |
| 205391 | Acidobacteria | Acidobacteria_Gp3 | 0.87 | 327,162 | 2,190 |
| 4378940 | Acidobacteria | Acidobacteria_Gp6 | 1 | 310,421 | 2,499 |
| 946250 | Verrucomicrobia | Spartobacteria | 1 | 300,544 | 2,386 |
| 3122801 | Acidobacteria | Acidobacteria_Gp1 | 0.97 | 273,493 | 2,107 |
| 4450676 | Proteobacteria | Alphaproteobacteria | 1 | 255,424 | 1,963 |
| 206514 | Proteobacteria | Alphaproteobacteria | 0.92 | 207,460 | 2,075 |
| 4463040 | Firmicutes | Clostridia | 0.94 | 204,483 | 2,096 |

### Single cell genomics

For single cell genomics, a soil sample was collected from 0-10 cm depth in a residential garden in Nobleboro, Maine (44° 5’48.10"N, 69°29’10.56"W) on May 5^th^, 2015. Approximately five grams of the sample were mixed with 30 mL sterile-filtered phosphate-saline buffer (PBS), vortexed for 30 s at maximum speed, and centrifuged for 30 s at 2,000 rpm. The obtained supernatant was diluted to below 10^5^ cell x mL^−1^ with PBS, pre-screened through a 40 μm mesh-size cell strainer (BD), and incubated with SYTO-9 DNA stain (5 μM; Invitrogen) for 10-60 min. The generation of single amplified genomes (SAGs) and their genomic sequencing were performed by the Bigelow Laboratory Single Cell Genomics Center (scgc.bigelow.org), as previously described(Stepanauskas *et al).* SAGs representing the “most-wanted list" were selected based on 16S rRNA gene BLAST alignments with greater than 97% similarity over at least 72 bp.

### Code and sequencing data availability

The analysis code used to generate the results is available from https://github.com/germs-lab/refsoil. All 14 single-cell sequencing data sets have been deposited at the NCBI. Sequencing data with NCBI accession identifier are listed in Table 2.

**Table 2.** Single-cell amplified genomes. Taxonomic classification of single-cell amplified genomes and the abundance of the most similar EMP OTU.

| NCBI ID | Phylum | Class | EMP ID in (Rideout et al., 2014) | Abundance | Relative Abundance (%) |
| --- | --- | --- | --- | --- | --- |
| LSTF00000000 | Proteobacteria | β-proteobacteria | New.52.CleanUp.ReferenceOTU25474 | 81,630 | 2.13E-02 |
| LSTI00000000 | Actinobacteria | Thermoleophilia | 551344 | 35,500 | 9.27E-03 |
| LSTA00000000 | Verrucomicrobia | Spartobacteria | New.54.CleanUp.ReferenceOTU60738 | 5,076 | 1.33E-03 |
| LSTC00000000 | Nitrospirae | Nitrospira | New.22.CleanUp.ReferenceOTU3952 | 4,337 | 1.13E-03 |
| LSTH00000000 | Planctomycetes | Planctomycetia | 521158 | 4,161 | 1.09E-03 |
| LSSZ00000000 | Planctomycetes | Planctomycetia | New.5.CleanUp.ReferenceOTU188938 | 2,434 | 6.36E-04 |
| LSTE00000000 | Proteobacteria | γ-proteobacteria | NA | NA | NA |
| LSTB00000000 | Proteobacteria | α-proteobacteria | New.52.CleanUp.ReferenceOTU4588 | 309 | 8.07E-05 |
| LSSY00000000 | Actinobacteria | Thermoleophilia | 904196 | 103 | 2.69E-05 |
| LSTG00000000 | Verrucomicrobia | Opitutae | New.5.CleanUp.ReferenceOTU291124 | 69 | 1.80E-05 |
| LSTD00000000 | Planctomycetes | Planctomycetia | New.59.CleanUp.ReferenceOTU1223798 | 44 | 1.15E-05 |
| LSTJ00000000 | Verrucomicrobia | Verrucomicrobiae | New.7.CleanUp.ReferenceOTU84651 | 11 | 2.87E-06 |
| LSTK00000000 | Acidobacteria | Acidobacteriia | New.59.CleanUp.ReferenceOTU999655 | 4 | 1.04E-06 |
| LSSX00000000 | Chloroflexi | TK17 | New.59.CleanUp.ReferenceOTU481621 | 2 | 5.22E-07 |

### Results

#### Phylogenetic and functional characterization of RefSoil

RefSoil is a manually curated database containing a total 922 genomes of soil-associated organisms, comprised of 888 bacteria and 34 archaea (Supplementary Table 1). The genomes within RefSoil (soil-associated) and RefSeq (all environments) reference databases were compared. Both genome databases contained similar dominant phyla, including Proteobacteria, Firmicutes, and Actinobacteria, comprising over 91% and 88% of genomes in RefSoil and RefSeq bacteria, respectively. RefSoil contained higher proportions of Armatimonadetes, Germmatimonadetes, Thermodesulfobacteria, Acidobacteria, Nitrospirae, and Chloroflexi than RefSeq, suggesting that these phyla may be enriched in the soil or underrepresented in the RefSeq database. A total of eleven RefSeq-associated phylums were absent from RefSoil indicating that these phyla may be absent or difficult to cultivate in soil environments (Supplementary Fig. 1). The 16S rRNA genes for RefSoil bacterial genomes were obtained from NCBI Genbank annotations and aligned to construct a phylogenetic tree representing RefSoil phylogenetic diversity (Fig. 1).

RefSoil bacterial and archaeal genomes were further annotated with the Rapid Annotation using Subsystem Technology (RAST, v 2.0(Aziz *et al.*, 2008)) (Supplementary Table 1), resulting in the annotation of a total of 1,811,233 RAST-associated genes. RAST annotations, unlike GenBank annotations, include classification into functional ontologies or subsystems, allowing for the comparison of broad functions. Overall, 78% of the bacterial and archaeal genes could be classified into functional subsystems (Supplementary Table 2). For genes associated with each subsystem, we evaluated the phylogenetic origins of RefSoil genes and compared the phlya distribution of annotated genes with those within the cumulative RefSoil database (Fig. 2). If the proportion of genes associated with a phylum within a functional subsystem was greater than its representation in RefSoil, we considered the representation of the phyla *enriched* in this function (Fig. 2, Supplementary Table 3). For example, we observed that in most subsystems (15 out of 26 subsystems), Proteobacteria-associated genes were enriched relative to their representation within RefSoil (56% of all RefSoil genes associated with Proteobacteria). Additionally, Actinobacteria, Crenarchaeota, and Proteobacteria genes were enriched in functions related to metabolism of aromatic compounds; Firmicutes and Proteobacteria genes were enriched in functions related to iron acquisition and metabolism; and function category dormancy and sporulation was enriched with Firmicutes genes only. These enrichments indicate that organisms from a small number of phyla may have advantages over other organisms in environments where these functions are important.

#### RefSoil compared to sequences originating from soil environments

Genomes in RefSoil represent cultivated strains originating from soils that have been isolated and often characterized in laboratory conditions. In order to estimate their natural abundance in soils, we compared RefSoil to amplicon sequencing datasets from global soil microbiomes in the Earth Microbiome Project (EMP)(Rideout *et al.*, 2014; Gilbert *et al.*, 2014). Amplicons (16S rRNA genes) from EMP datasets were clustered at 97% sequence similarity to determine representative operational taxonomic units (OTUs)(Rideout *et al.*, 2014). We selected only EMP samples that were associated with soil samples, resulting in a total of 3,035 samples. Within these soil samples, we observed that the majority of OTUs were rare (e.g., only observed in a few samples), with 76% of OTUs observed in less than ten soil samples, and 1% of OTUs representing 81% of total abundance in EMP.

Comparing RefSoil 16S rRNA genes to EMP amplicons, we observed that the majority of RefSoil 16S rRNA sequences were highly similar to EMP amplicons (over 87% of RefSoil bacterial and archaeal 16s rRNA genes shared greater than 97% similarity with a minimum 72 bp alignment length). In contrast, 99% (2,442,432 of 2,476,795) of EMP amplicons did not share high similarity (greater than 97% similarity) to RefSoil genes, suggesting that EMP soil samples contain much higher diversity than represented within RefSoil. While RefSoil genomes represent soil microbes that have been well-studied in the laboratory, the EMP amplicons represent microbes that have been observed in soils around the world but not necessarily well-characterized (e.g., limited to the observation of the presence of 16S rRNA gene sequences).

#### Soil microbiomes associated with different soil types

In 1975, a soil taxonomy was developed by the United States Department of Agriculture (USDA) and the National Resources Conservation Service, which separates soils into twelve orders that are based on their physical, chemical, or biological properties(Soil Survay Staff 1999, 1999). Despite the availability of this classification, it is rarely incorporated into soil microbiome surveys. Using RefSoil and estimated abundances from similar EMP OTUs, we evaluated microbial distribution in various soil orders. We obtained GPS data from EMP soil samples originating from the United States that allowed us to obtain the soil classification. Within these EMP samples, the most represented soil orders included Mollisols (58%, grassland fertile soils) and Alfisols (37%, fertile soils typically under forest vegetation) (Supplementary Table 4). The abundance of OTUs similar to RefSoil genes was used to estimate the membership of well-characterized isolates in these soil samples (Supplementary Fig. 2). Mollisols, Alfisols and Vertisols (soils with high clay content with pronounced changes in moisture) were associated with the most RefSoil representatives, while Gelisols (cold climate soils), Ultisols (soils with low cation exchange), and sand/rock/ice contained the least (Supplementary Table 4).

#### Identification of the most beneficial future targets for RefSoil

EMP OTUs that do not share high similarity with 16S rRNA genes from RefSoil represent current knowledge gaps for which we lack cultivated isolates. These gaps were visually identified by overlaying the presence of highly similar relatives in RefSoil (similarity >97%) to branches in a phylogenetic tree of soil-associated EMP amplicons (Fig. 3). We evaluated the presence of OTUs in EMP libraries based on their presence in EMP samples (frequency) and their cumulative observed abundance. We identified the most observed and abundant EMP OTUs that did not share high similarity to Refsoil to generate a “RefSoils most wanted list” of targets that, if isolated or sequenced, could provide information on prevalent but uncharacterized lineages (Table 1, Supplementary Table 5-6). OTUs sharing similarity to Verrucomicrobia (8 OTUs) and Acidobacteria (6 OTUs) were among the most abundant and frequently observed EMP OTUs that are not currently represented in RefSoil (Table 1). These targets, observed in nature but not in our database, would be ideal to isolate and characterize to better understand the soil biodiversity.

#### Single cell genomics in soil

Sequencing-based approaches provide an alternative to accessing the genomes of soil organisms without cultivation. Previous efforts have used assembly of genomes from metagenomes(Labonté *et al.*, 2015; Martijn *et al.*, 2015; Field *et al.*, 2014) and single cell genomics(Gawad *et al.*, 2016; Lasken, 2012; Stepanauskas, 2012) to obtain genomic blueprints of yet uncultured microbial groups. To evaluate the effectiveness of single cell genomics on soil communities, we performed a pilot-scale experiment on a residential garden soil in Maine. The 16S rRNA gene was successfully recovered from 109 of the 317 single amplified genomes (SAGs). This 34% 16S rRNA gene recovery rate is comparable to single cell genomics studies in marine, freshwater and other environments(Martinez-Garcia *et al.*, 2011; Rinke *et al.*, 2013; Swan *et al.*, 2011). The 16S rRNA genes of 14 of these SAGs, belonging to Proteobacteria, Actinobacteria, Nitrospirae, Verrucomicrobia, Planktomycetes, Acidobacteria and Chloroflexi, were selected based on their lack of representation within RefSoil. Genomic sequencing of those SAGs resulted in a cumulative assembly of 23 Mbp (Table 2, Supplementary Table 7). We estimated the abundance of OTUs similar to single cell 16S rRNA genes to range from 5E-7 to 2E-2% of EMP OTU abundances. These abundances are very low but comparable to the average and median relative abundance of OTUs within EMP (4E-3% and 1E-6%, respectively). If these draft genomes were added to RefSoil, these 14 SAGs would increase RefSoils representation of EMP amplicons by 7% by abundance.

### Discussion

Advances in sequencing techniques for utilizing culture-independent approaches have created new opportunities for understanding soil microbiology and its impact on soil health, stability, and management. We provide RefSoil as a community genomic resource to provide high-quality curation of organisms that originate in soil studies, allowing us to more accurately annotate soil sequencing datasets. This database is an important initial effort to provide improved soil references, and we acknowledge that it is far from a complete representation of soils biodiversity. In fact, compared to EMP soils, RefSoil represents only 2% of observed EMP OTUs and 10% by abundance of the observed soil OTUs, highlighting the magnitude of soil biodiversity and the limitations of our current database based on cultivated representatives. These results are consistent with a recent effort demonstrating that we are very limited in our knowledge of not only soil but lifes diversity and advocating that much can be learned by including uncultivated organisms (Hug *et al.*, 2016). We show here how RefSoil can be used to evaluate which organisms to target that would most effectively increase our knowledge of soil biodiversity. For example, if genome references were available for the top most wanted organisms identified in this effort (Table 1), we could expand RefSoils representation of EMP soils by 58% by abundance.

Notably, many of these targets have previously been observed to be recalcitrant to cultivation with standard laboratory media, resulting in their absence in current genome databases. Acidobacteria, for example, is known to be slow growing and difficult to cultivate (Nunes da Rocha *et al.*, 2009) while also observed to be highly abundant in soil (33% of EMP amplicons by abundance). Another fastidious bacteria, Verrucomicrobia(Bergmann *et al.*, 2011), were also observed to be highly abundant (12.5%) in EMP but not well represented in RefSoil (two of 888 bacterial genomes). Despite their absence from cultivated isolates, both Acidobacteria and Verrucomicrobia have been observed to be critical for nutrient cycling in soils (Nunes da Rocha *et al.*, 2009; Ward *et al.*, 2009; Fierer *et al.*, 2013; Martinez-Garcia *et al.*, 2012).

Novel isolation and culturing techniques will help us to access these previously difficult to grow bacteria and will also be complemented by emerging sequencing technologies. In particular, single-cell genomics hold great promise to provide genomic characterization of lineages that are difficult to culture (Gawad *et al.*, 2016; Lasken, 2012; Stepanauskas, 2012). In our pilot experiment, we demonstrate, for the first time, that single-cell genomics is applicable on soil samples and is well suited to recover the genomic information from abundant but yet uncultured taxonomic groups. The 14 sequenced SAGs have significantly increased the extent to which RefSoil represents the predominant soil lineages from a single sample. Much larger single cell genomics projects are feasible and have been employed in prior studies of other environments(Kashtan *et al.*, 2014; Rinke *et al.*, 2013; Swan *et al.*, 2013). The continued, rapid improvements in this technology are likely to lead to further scalability, offering a practical means to fill the existing gaps in the RefSoil database and biodiversity more broadly.

The RefSoil database is also a tool that allows us to characterize currently known soil bacteria phylogeny and functions. This database spans 24 phyla of bacteria and archaea. While genes related to microbial growth and reproduction (e.g., DNA, RNA, and protein metabolism) originate from diverse phyla, key functions related to metabolism of aromatic compounds, iron metabolism, and dormancy and sporulation were observed to be enriched from only a few RefSoil phyla, suggesting that these phyla may have specialized functions within soil communities. Specifically, Proteobacteria-associated genes were enriched in functions related to motility, chemotaxis, and membrane transportation (Figure 2), suggesting that Proteobacteria are likely to be efficient at acquiring readily available nutrients and elements for growth(Fierer *et al.*, 2007). Genes related to Proteobacteria and Actinobacteria were also found dominant among RefSoil genomes in subsystems related to the metabolism of aromatic compounds. This observation is consistent with the association of Proteobacteria and humic substances utilization and the contribution of Actinobacteria to plant material degradation(Godden *et al.*, 1992; Fuchs *et al.*, 2011).

By comparing RefSoil to other databases, we identified biases for specific phyla in RefSoil; in particular, Firmicutes are observed frequently in RefSoil but were not observed to be highly abundant in soil environments (5.7% of all EMP amplicons). Firmicutes have been well-studied as pathogens(Buffie and Pamer, 2013; Kamada *et al.*, 2013; Rupnik *et al.*, 2009), likely biasing their representation in our databases and consequently their annotations in soil studies. A key advantage to the development of the RefSoil database is the opportunity to evaluate these biases and to identify targets for future curation that would create a more representative database resource for the soil. To this end, we evaluated the representation of RefSoil in various soils.

Consistent with previous observations that microbial community composition is correlated with soil environments(Fierer *et al.*, 2012; Fierer and Jackson, 2006), we observed specific RefSoil membership associated with soil taxonomy. Not all soil taxonomic orders were equally represented in RefSoil, with the most genomes associated with Mollisols, or grassland soils. This bias is likely due to increased research in these soils due to their importance for agriculture and productivity. In contrast, for Gellisol permafrost soils, which cover over 20% of Earths terrestrial surface(Koven *et al.*, 2011) and are a significant carbon source from our biosphere to the atmosphere, we observe only a small fraction of known soil microbes are present. Previous studies suggest that high levels of novelty can be observed in permafrost microbiomes(Koven *et al.*, 2011; Mackelprang *et al.*, 2011), suggesting that these soils could significantly benefit from improved references.

Another advantage to the RefSoil database is that it represents genomes for which strains should currently be available and for which we have high-quality genomes. As a consequence, genomes within RefSoil could help to inform and design soil microbiology experiments. For example, it is known that nitrogen cycling genes are abundant in agricultural soils (Xue *et al.*, 2013; Philippot *et al.*, 2007; Long *et al.*, 2012). A mock community of isolates known for participating in nitrogen cycling could be generated and proportionally designed to mimic soil conditions using RefSoil: these strains might include microorganisms related to *Streptomyces venezuelae* ATCC 10712 (assimilatory nitrate reductase), *Bacillus anthracis* (nitric oxide reductase), *Pseudomonas brassicacearum* (nitrous oxide reductase), *Halomonas elongata* DSM 2581 (ammonia monooxygenase)*, Pseudomonas fluorescens* SBW25 (ammonia monooxygenase), and *Pleurocapsa sp.* PCC 7327 (nitrogen fixation). Combining RefSoil and available amplicon datasets, one could estimate the proportions of strains as observed in soil samples.

In conclusion, RefSoil is an important first step towards building a more comprehensive, well-curated database. Though it is far from complete, we have been able to use RefSoil as a tool to identify key phyla that represent gaps in known soil diversity and underrepresented phyla. Going forward, this reference will be expanded as isolation and cultivation-independent technologies continue to improve.

## Acknowledgments

We are grateful for the support of the NSF Terragenome International Soil Metagenome Sequencing Consortium for providing a collaborative workshop with the soil community for discussions to improve this project. This material is based upon work supported by the U.S. Department of Energy, Office of Science, Office of Biological and Environmental Research, under Award Number SC0010775.

**Contributions** E.C., A.G., A.H., and J.T. curated Refsoil; J.C. and A.H. collected the genome information to build the database; J.C, J.F., A.H., R.W, and F.Y., characterized and compared RefSoil to EMP datasets; B.G. provided soil orders for EMP samples; R.S. performed single cell genomics; J.T., K.H. and A. H. supervised the work; J.C, F.Y., and A.H. wrote the manuscript with contributions from all other authors.

## Figure Legends

### Figure 1. Phylogenetic tree of RefSoil

Phylogenetic tree of aligned 16S rRNA genes originating from RefSoil bacterial genomes. A: Acidobacteria, B: Actinobacteria, C: Aquificae, D: Armatimonadetes, E: Bacteroidetes, F: Chlamydiae, G: Chlorobi, H: Chloroflexi, J: Cyanobacteria, K: Deferribacteres, L: Deinococcus-Thermus, N: Firmicutes, O: Fusobacteria, P: Gemmatimonadetes, Q: Nitrospirae, R: Planctomycetes, S: Proteobacteria, T: Spirochaetes, U: Synergistetes, V: Tenericutes, W: Thermotogae, X: Verrucomicrobia.

### Figure 2. Functional analysis

The distribution of phylogenetic origins associated with RefSoil functional subsystems as annotated by RAST (panel A); the overall phylogenetic distribution of all genes in RefSoil (panel B).

### Figure 3. Phylogenetic tree of EMP OTUs clustered by taxonomy

Ring I (green) represents the cumulative log-scaled abundance of OTUs in EMP soil samples. Ring II (red) represents EMP OTUs that share greater than 97% gene similarity (to RefSoil 16S rRNA genes; ring III (blue) indicates that these 16S rRNA genes shared similarity to sorted cells that were selected for single cell genomics. A: Acidobacteria, B: Actinobacteria, C: Aquificae, D: Armatimonadetes, E: Bacteroidetes, F: Chlamydiae, G: Chlorobi, H: Chloroflexi, I: Crenarchaeota, K: Deferribacteres, L: Deinococcus-Thermus, M: Euryarchaeota, N: Firmicutes, O: Fusobacteria, P: Gemmatimonadetes, Q: Nitrospirae, R: Planctomycetes, S: Proteobacteria, T: Spirochaetes, U: Synergistetes, V: Tenericutes, W: Thermotogae, X: Verrucomicrobia, Y: Cyanobacteria/Chloroplast.

## Table Legends

### Table 1. RefSoil’s most wanted OTUs

RefSoils most wanted OTUs based on observed frequency and abundance in EMP soil samples. Taxonomy for OTUs are assigned by RDP classifier(Wang *et al.*, 2007). *:(Rideout *et al.*, 2014).

### Table 2. Single-cell amplified genomes

Taxonomic classification of single-cell amplified genomes and the abundance of the most similar EMP OTU. *:(Rideout *et al.*, 2014).

## Supplementary Description

### Supplementary Figure 1. The abundance distribution of phyla

The abundance distribution of phyla (log(abundance) + 1) present in the RefSoil database compared to NCBIs RefSeq (A) and phyla that are enriched in RefSoil compared to RefSeq (B).

### Supplementary Figure 2. Average relative abundance for various soil orders

Average relative abundance of EMP OTUs (sharing similarity with RefSoil genes) for various soil orders as classified by NRCS Soil Taxonomy.

### Supplementary Table 1. The RefSoil Database

### Supplementary Table 2. RefSoil genes in RAST subsystem functions

Total number of RefSoil genes observed in RAST subsystem functions.

### Supplementary Table 3. Subsystem level functions in RefSoil

The number of unique phyla and the number of enriched phyla associated with encoded subsystem level functions in RefSoil. (Bold italicized: enriched with genes associated with three or less phyla)

### Supplementary Table 4. Abundance of EMP OTUs in soil order

The total abundance of EMP OTUs by soil order in soil samples originating in the United States.

### Supplementary Table 5.100 most abundant OTUs

100 most abundant OTUs in EMP soil samples

### Supplementary Table 6.100 most frequent OTUs

100 most frequent OTUs in EMP soil samples

### Supplementary Table 7. Single cell genomic properties

